# Junction-based lamellipodia drive endothelial cell rearrangements *in vivo* via a VE-cadherin/F-actin based oscillatory ratchet mechanism

**DOI:** 10.1101/212522

**Authors:** Ilkka Paatero, Loïc Sauteur, Minkyoung Lee, Anne K. Lagendijk, Daniel Heutschi, Dimitri Bieli, Benjamin M. Hogan, Markus Affolter, Heinz-Georg Belting

## Abstract

Angiogenesis and vascular remodeling are driven by a wide range of endothelial cell behaviors, such as cell divisions, cell movements, cell shape and polarity changes. To decipher the cellular and molecular mechanism of cell movements, we have analyzed the dynamics of different junctional components during blood vessel anastomosis *in vivo*. We show that endothelial cell movements are associated with oscillating lamellipodia-like structures, which are orientated in the direction of these movements. These structures emerge from endothelial cell junctions and we thus call them junction-based lamellipodia (JBL). High-resolution time-lapse imaging shows that JBL are formed by F-actin based protrusions at the front end of moving cells. These protrusions also contain diffusely distributed VE-cadherin, whereas the junctional protein ZO-1 (Zona occludens 1) remains at the junction. Subsequently, a new junction is formed at the front of the JBL and the proximal junction is pulled towards the newly established distal junction. JBL function is highly dependent on F-actin dynamics. Inhibition of F-actin polymerization prevents JBL formation, whereas Rac-1 inhibition interferes with JBL oscillations. Both interventions disrupt endothelial junction formation and cell elongation. To examine the role of VE-cadherin (encoded by *cdh5 gene*) in this process, we generated a targeted mutation in VE-cadherin gene (*cdh5*^*ubs25*^), which prevents VE-cad/F-actin interaction. Although homozygous *ve-cadherin* mutants form JBL, these JBL are less dynamic and do not promote endothelial cell elongation. Taken together, our observations suggest a novel oscillating ratchet-like mechanism, which is used by endothelial cells to move along or over each other and thus provides the physical means for cell rearrangements.

## Introduction

Organ morphogenesis is driven by a wealth of tightly orchestrated cellular behaviors, which ensure proper organ assembly and function. The cardiovascular system is one of the most ramified vertebrate organs and is characterized by an extraordinary plasticity. It forms during early embryonic development, and it expands and remodels to adapt to the needs of the growing embryo. In adult life, this plasticity allows flexible responses, for example, during inflammation and wound healing ^1,2^.

At the cellular level, blood vessel morphogenesis and remodeling are accomplished by endothelial cell behaviors including cell migration, cell rearrangement and cell shape changes ^3–5^. This repertoire of dynamic behaviors allows endothelial cells to rapidly respond to different contextual cues, for example during angiogenic sprouting, anastomosis, diapedesis or regeneration. In particular, it has been shown that endothelial cells are very motile, not only during sprouting, but also within established vessels, where they migrate against the blood flow ^6,7^.

Endothelial cell migration has been extensively studied in different *in vivo* and *in vitro* systems mainly focusing on angiogenic tip cell behavior and the interaction of endothelial cells with the extracellular matrix (ECM) ^8,9^. However, endothelial cells can also shuffle positions within an angiogenic sprout ^10^ and these cellular rearrangements require the junctional adhesion protein VE-cadherin/CDH5 ^11–13^. Moreover, *in vivo* analyses in avian and fish embryos have shown that endothelial cells can migrate within patent blood vessels emphasizing that regulation of endothelial cell adhesion and motility is critical during vascular remodeling processes ^6,7,14^.

Although many aspects of sprouting angiogenesis and vascular remodeling rely on endothelial cell interactions ^3^, the exact role of endothelial cell junctions (and in particular that of VE-cad) in these processes is not well understood. Indeed, rather than supporting an active function for VE-cad in dynamic cell behaviors, most studies point to a restrictive or permissive role, consistent with the maintenance of endothelial integrity ^15–17^. On the other hand, the observation that loss of VE-cad function can inhibit cell rearrangements suggests an active contribution to this process ^12,13^.

To decipher the cellular and molecular mechanisms, which enable cells to move within the endothelium, we have focused on the process of anastomosis during the formation of the dorsal longitudinal anastomotic vessel (DLAV) in the zebrafish embryo by high-resolution time-lapse microscopy. This process occurs in a relatively stereotypical manner and involves a convergence movement of endothelial cells, which is illustrated by extensive cell junction elongation. Ultimately, this process alters tube architecture and converts unicellular vessels to multicellular vessels. By *in vivo* time-lapse imaging of several junctional components and pharmacological interference with F-actin dynamics, we are able to describe a novel actin-based mechanism, which allows endothelial cells to move along each other while maintaining junctional integrity. In particular, we describe a novel rearrangement mechanism, which is initiated by junction-based-lamellipodia (JBL) leading to the formation of distal, VE-cad based attachment sites, which in turn serve as an anchor point for junction elongation. We propose that the oscillating behavior of JBL, which depends on F-actin polymerization as well as contractility, provides a general mechanism of endothelial cell movement during blood vessel formation and vascular remodeling.

## Results

### Changes of vessel architecture during blood vessel formation

Blood vessel formation is associated with prominent cell shape changes and cell rearrangements. The DLAV presents a well-defined model to analyze how a wide repertoire of endothelial cell activities leads to the formation of a new blood vessel, starting with establishment of an interendothelial contact point, followed by the formation of a continuous luminal surface and the transformation from a unicellular to a multicellular tubular architecture. Unicellular and multicellular tubes are easily discerned by junctional patterns: whereas unicellular tubes display isolated rings separated by segments without any junction, multicellular tubes have a continuous network of multiple junctions along their longitudinal axis. To gain more insight into this transformation process, we generated a reporter line expressing a full-length VE-cadherin fluorescent protein fusion (VE-cad-Venus) (Fig. 1A-D, Supplemental Movie 1) and performed *in vivo* time-lapse experiments between 27 and 40 hours post fertilization (hpf). These experiments showed that most DLAVs were initially unicellular tubes, and that the majority (69%, n=26 (8 embryos)) of DLAV segments were transformed to a multicellular configuration before 40 hpf (Fig. 1A-D, Supplemental Fig. 1A). The transformation from the initial unicellular contact to a multicellular vessel, with a continuous cell-cell junction network, took several hours (median 190 min, segments n=14 (8 embryos), Supplemental Fig.1B), with high variability between individual segments. During this time window, the endothelial cell-cell junctions expanded extensively from initial spot-like structures to elongated junctions covering the entire DLAV segment. However, movement of the junctions was also seen in perfused vessels. The cellular rearrangements are thus occurring both in nascent non-lumenized vessels and also inflated, perfused vessels.

**Figure 1.**
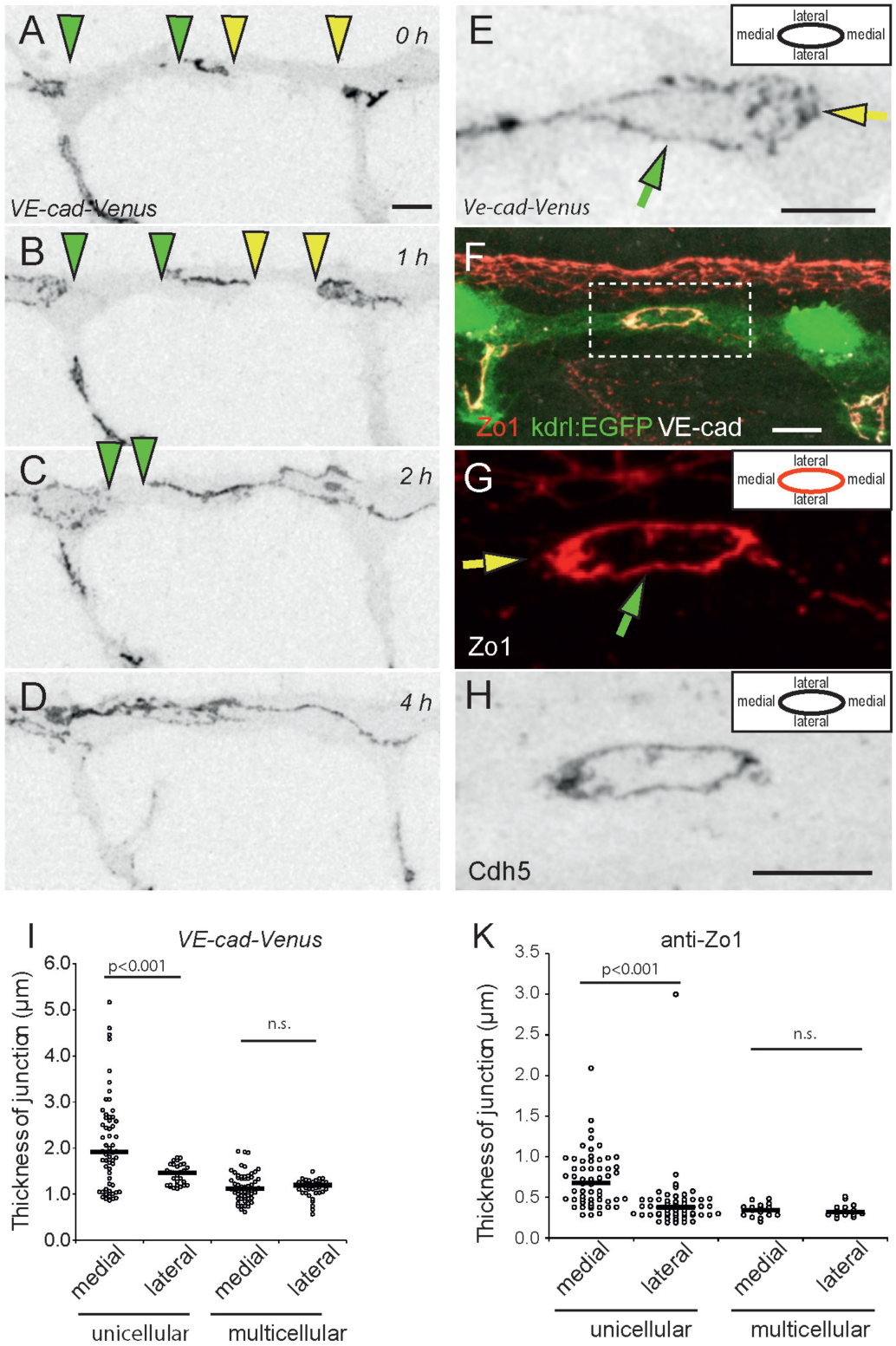
Polarized thickness in remodeling junctions. A-D) Still pictures of a time-lapse movie (Supplemental movie 1) showing EC junctions of a *Tg(BAC(cdh5:cdh5-ts)) embryo*, which expresses VE-cad-Venus fusion protein, during transition from unicellular to multicellular vessel architecture during DLAV formation, in inversed contrast starting around 28 hpf. The edges of the junctional gaps in the unicellular vessels are marked with green and yellow triangles. E) Close-up image of VE-cad-Venus embryos. The diffuse appearance of the medial junction is marked by a yellow arrow, the more condensed lateral junction by a green arrow. F-H) Whole-mount immunofluorescence staining of the DLAV of 28-30hpf kdrl:EGFP (Tg(*kdrl:EGFP*^*s843*^)) embryos using anti-ZO1 and anti-VE-cad antibodies. In G the yellow arrow points to medial junctional domain and green arrow to lateral junctional domain. I) Quantitation of junctional thickness measurements. n= 44 for unicellular junctions and n=48 for multicellular junctions from 5 VE-cad-Venus *(Tg(BAC(cdh5:cdh5-ts*))) embryos. Non-parametric Kruskal-Wallis test was used. K) Quantitation of the junctional thickness measurements from immunostainings of Tg(*kdrl:EGFP*^*s843*^) embryos. n=58 for unicellular junctions and n=17 for multicellular junctions from 8 embryos. Non-parametric Kruskal-Wallis test was used. Scale bars 10µm.

### The thickness of remodeling junctions is polarized

When we analyzed cell-cell junctions in the DLAV of VE-cad-Venus embryos in more detail, we observed that the junctions were not uniform in thickness along their circumference in unicellular vessels. In medial regions, the junctions were thicker and showed a more diffuse pattern than on the lateral sides (Fig. 1E and I), coinciding with the general direction of endothelial cell movements during anastomosis. In contrast, we did not observe such junctional polarity in multicellular vessels (Fig. 1I). We confirmed these observations by immunostainings for the junctional proteins VE-cadherin and Zo-1, which showed that in the newly formed junctions in unicellular configuration, the medial junctional domains were consistently thicker than the lateral domains (Fig. 1F, G, H and K). Again, this polarity of junctional thickness was not seen in vessel areas of more mature multicellular architecture (Fig. 1K).

### Remodeling junctions form dynamic junction-based lamellipodia

To gain insight into the nature of this junctional polarity, we performed live-imaging experiments on Cdh5-Venus expressing transgenic embryos at high temporal resolution. Here, we observed that the polarized junctional thickenings are formed by dynamic lamellipodia-like protrusions (Fig. 2A and B, Supplemental Movie 2). In addition, we used a F-actin visualizing EGFP-UCHD transgenic fish line in a similar setup (Figure 3A and B, Supplemental Movie 3). Remarkably both, F-actin and VE-cadherin, showed similar oscillatory dynamics with a median duration of 6 minutes (Fig. 2E). Moreover, the protrusions were oriented along the vessel axis (Fig. 2F), which is consistent with the increased junctional thickness of medial junctional domains.

**Figure 2.**
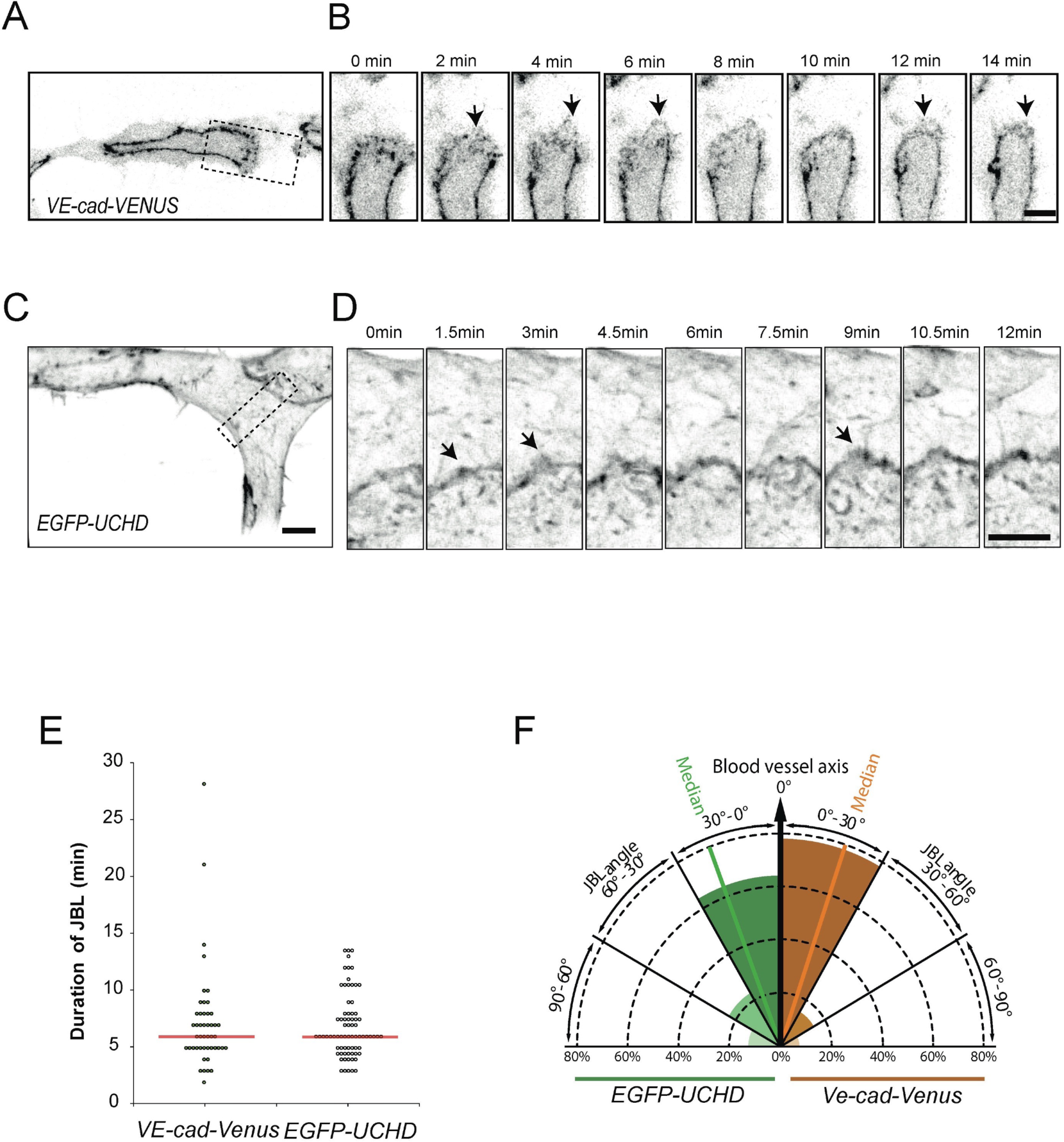
Oscillatory dynamics VE-cad and F-actin in junction-based lamellipodia. A-B) Still images from a movie (Supplemental movie 2) of a VE-cad-Venus expressing embryo (*Tg(BAC(cdh5:cdh5-ts*))), showing the DLAV at 30 hpf in inversed contrast. B) Magnification of the inset in A. Arrows point to JBL. C-D) Still images from a movie (Supplemental movie 3) of a EGFP-UCHD expressing embryo (*Tg(fli1a:Gal4ff*^*ubs3*^, *UAS:EGFP-UCHD*^*ubs18*^)) showing the DLAV at 30hpf in inversed contrast. D) Magnification of the inset in C. Arrows point to JBL. E) Scatter plot of quantitation of the duration of the JBL with the VE-cad-Venus transgene (n=48 in 6 embryos) and EGFP-UCHD movies (n=74 in 6 embryos), respectively, red line represents the median. F) Quantitation of JBL angle in the DLAV with respect to the blood vessel axis (0°) using the EGFP-UCHD transgene (n=103 from 6 embryos) or Cdh5-Venus transgene (n= 41 from 5 embryos). Scale bars 5 µm.

**Figure 3.**
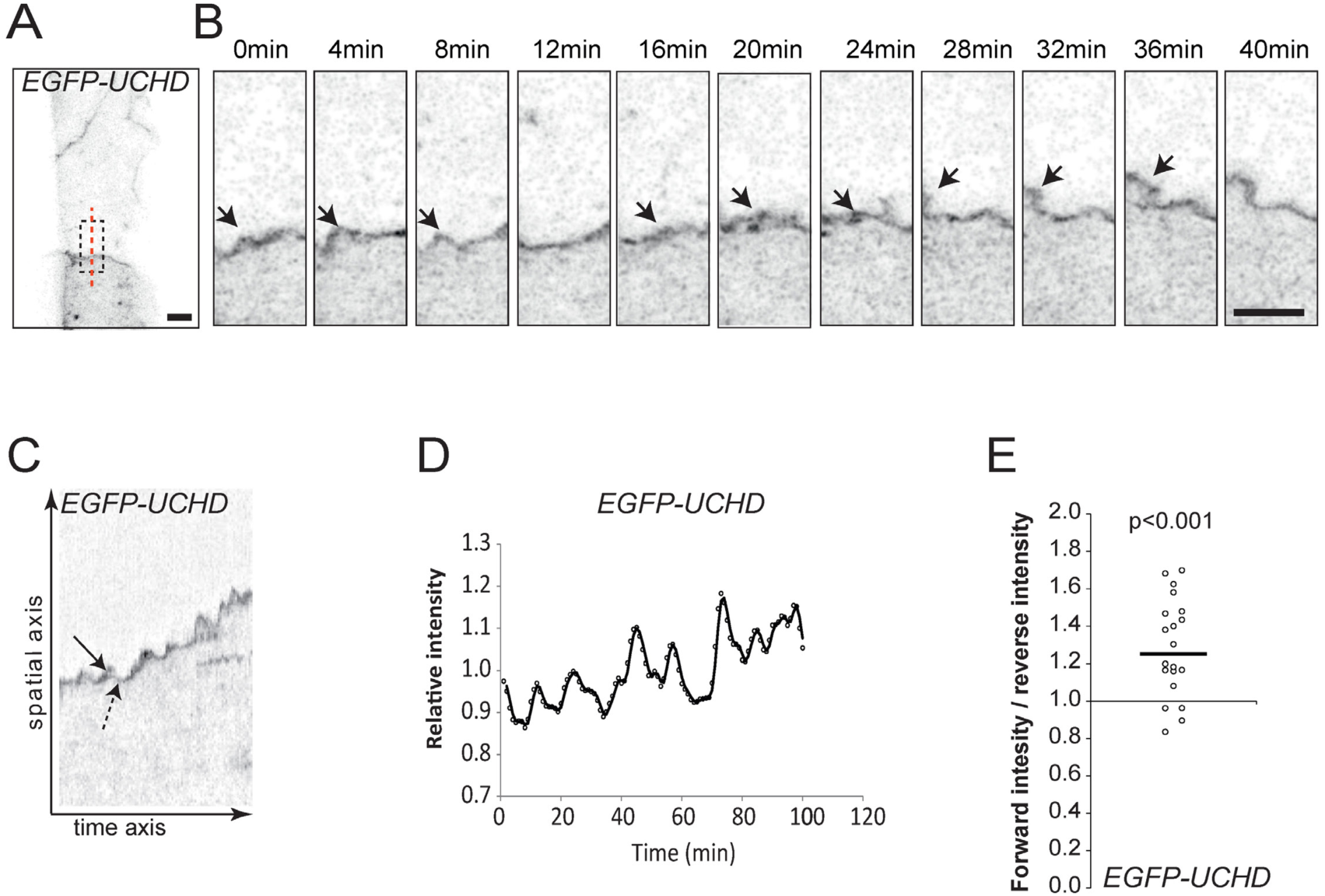
Oscillatory F-actin dynamics during remodelling of cell-cell junctions. A-B) Still images from a supplemental movie 4 showing JBL formation in the dorsal aorta of an EGFP-UCHD expressing 2dpf embryo (*Tg(fli1a:Gal4ff*^*ubs3*^; *UAS:EGFP-UCHD*^*ubs18*^)), shown in inversed contrast. B) Magnification of the inset in A). Red dashed line indicates the site for kymograph in C). Arrows point to a JBL, seen as a local thickening of the junction. C) Kymograph across the junction. Solid arrow denotes forward movement and dashed arrow backward movement of the junction. D) Intensity plotting of an EGFP-UCHD JBL kymograph. E) Scatter plot of the relative EGFP-UCHD intensity during forward and backward movements (n=20 events, 4 movies). EGFP-UCHD intensity value in a forward movement was divided with intensity value during subsequent reverse movement. Non-parametric one sample Wilcoxon signed rank test was used as statistical test. Scale bars 5 µm.

To test whether this dynamic junctional behavior is restricted to the process of anastomosis or is a more general behavior, we further analyzed the F-actin fluctuations during junctional remodeling in the dorsal aorta. Here, we found that the relative intensity of EGFP-UCHD was increased at the site of forming protrusions (Fig. 3A, Supplemental Movie 4), indicating recruitment of additional F-actin. We further analyzed the dynamic behavior of junctional F-actin using kymographs (Fig. 3C, D) and found that the fluctuations in intensity occurred in rather rhythmic patterns, and at the same junctional site, protrusions were generated repeatedly with in regular intervals, indicative for oscillations in F-actin intensity in the remodeling junction.

The polarized occurrence and directionality of protrusions along the direction of vessel growth suggests that they are involved in endothelial cell movements. To address this, we analyzed the potential association between local junctional movements and the occurrence of F-actin protrusions in the dorsal aorta (Fig. 3C and D). Analysis of F-actin intensities showed that higher intensities were associated with local forward movement of junctions than with reverse movement (Fig. 3E).

Taken together, we observe an F-actin based protrusive endothelial behavior, which occurs during junctional remodeling *in vivo*. Because of their similarity to “classical” lamellipodia, their oscillating behavior and structural connection with endothelial cell junctions, we call these protrusions junction-based lamellipodia or JBL.

### JBL form at the front end of elongating junctions

Blood vessel anastomosis is driven by the convergence movement of two tip cells and is associated with an elongation of their mutual cell junction. The formation of JBL at the junctional poles suggested that these dynamic structures may generate tractive forces, which contribute to junction elongation. However, cell junctions demarcate the interface between two cells and our above analyses did not differentiate, whether cells form JBL at their respective junctional front or rear ends or both. To analyze the contributions of individual cells to F-actin protrusions, we generated a transgenic zebrafish line expressing a photoconvertible mCLAV-UCHD in endothelial cells. For mosaic analysis, we photoconverted UCHD-mCLAV in single SeA sprouts prior to contact initiation and recorded the behavior of the junctional F-actin during anastomosis and formation of the DLAV (Fig 4A). *In vivo* photoconversion resulted in efficient green-to-red conversion (Fig. 4B), allowing the analysis of differentially labeled F-actin during DLAV formation (Fig. 4C and D). This analysis confirmed that the JBL are F-actin containing protrusions. Furthermore, each intercalating cell formed a single JBL at the front end of the junction with respect to cell movement (green JBL over red cell, or vice versa, 24 out of 28 events, p < 0.001). To separate the temporal order of cell interactions more clearly, we compared the dynamic distribution of F-actin at a single JBL in both contacting cells: the “donor cell”, which generates the JBL, and the “recipient cell”, which provides the adhesive substrate for the JBL. The JBL originated as a F-actin protrusion at the front end of the “donor” cell (Fig. 4D, red channel). This was associated with accumulation of F-actin to the site within the (non-photoconverted) recipient cell (Fig. 4D, green channel) (this occurred in 9 out of 10 analyzed events). These results show that cell interactions at JBL occur in a two-step process: an initial actin F-accumulation takes place at the front end of the elongating junction, which subsequently induces a secondary F-actin accumulation in the underlying cell.

**Figure 4.**
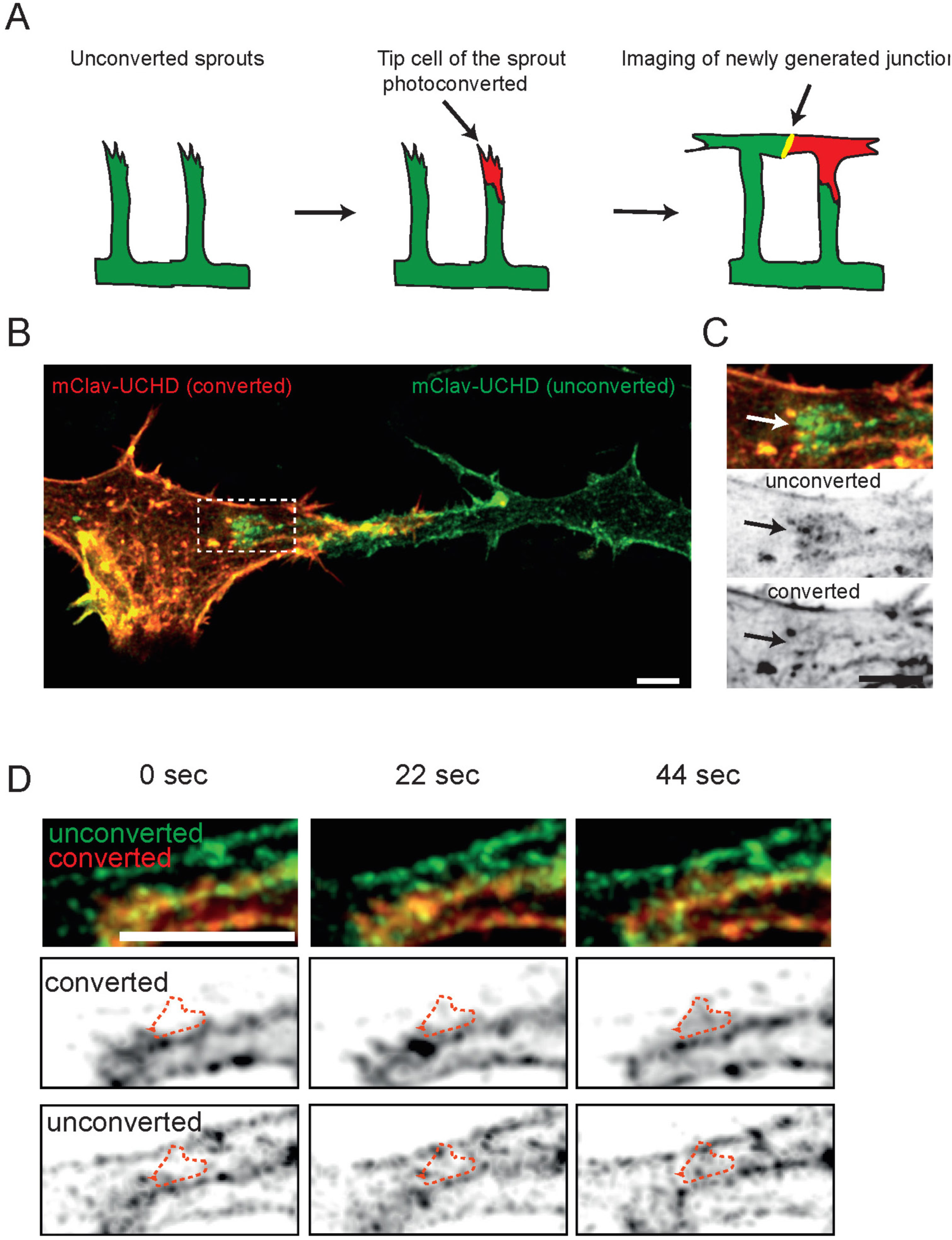
JBL formation at the distal tip of the junction during DLAV anastomosis. A) Schematic representation of the mClav2-UCHD photoconversion experiment. B) Image of photoconverted and unconverted mClav2-UCHD cells in the DLAV of an *Tg(fli1a:Gal4ff*^*ubs3*^; *UAS:mClav2-UCHD*^*ubs27*^) embryo, at 32 hpf. C) is a close up of the inset in B). Arrows point to differentially labelled JBL. D) Time lapse analysis of a single JBL formation between unconverted and photoconverted endothelial cells expressing mClav2-UCHD (*Tg(fli1a:Gal4ff*^*ubs3*^;*uas:mClav2-UCHD*^*ubs27*^)). Red-dashed lined outlines the emerging JBL. Scale bars 5 µm.

### F-actin protrusions precede junctional movements

Since the above observations suggest a step-wise mechanism of cell-cell interaction during JBL function, we set out to explore the spatio-temporal relationship between F-actin dynamics and the dynamics of other junctional components. To this end we generated transgenic fish lines expressing red F-actin (mRuby2-UCHD), which allows a direct comparison with other fluorescently labeled junctional components (e.g. EGFP-ZO1 and VE-cad-Venus) (Fig. 5A-B and C-D, respectively; Supplemental movies 5 and 6). Both VE-cad-Venus and EGFP-ZO1 followed the junctional F-actin front (11 and 9 movies analyzed, respectively); however, a different distribution pattern was observed during JBL formation. VE-cadherin localized diffusely at the front, largely overlapping with the F-actin protrusions (Fig. 5B, 60 to 120 sec.). In contrast, EGFP-ZO1 showed a more defined localization and initially remained associated with the junction at the proximal end of the protrusion (Fig. 5D 0 to 36 sec.). However, at later time points (Fig. 5D, 72 sec.), we observed EGFP-ZO1 accumulation also at the front edge of the JBL, indicating the formation of a new junction at this site (Fig. 5D).

**Figure 5.**
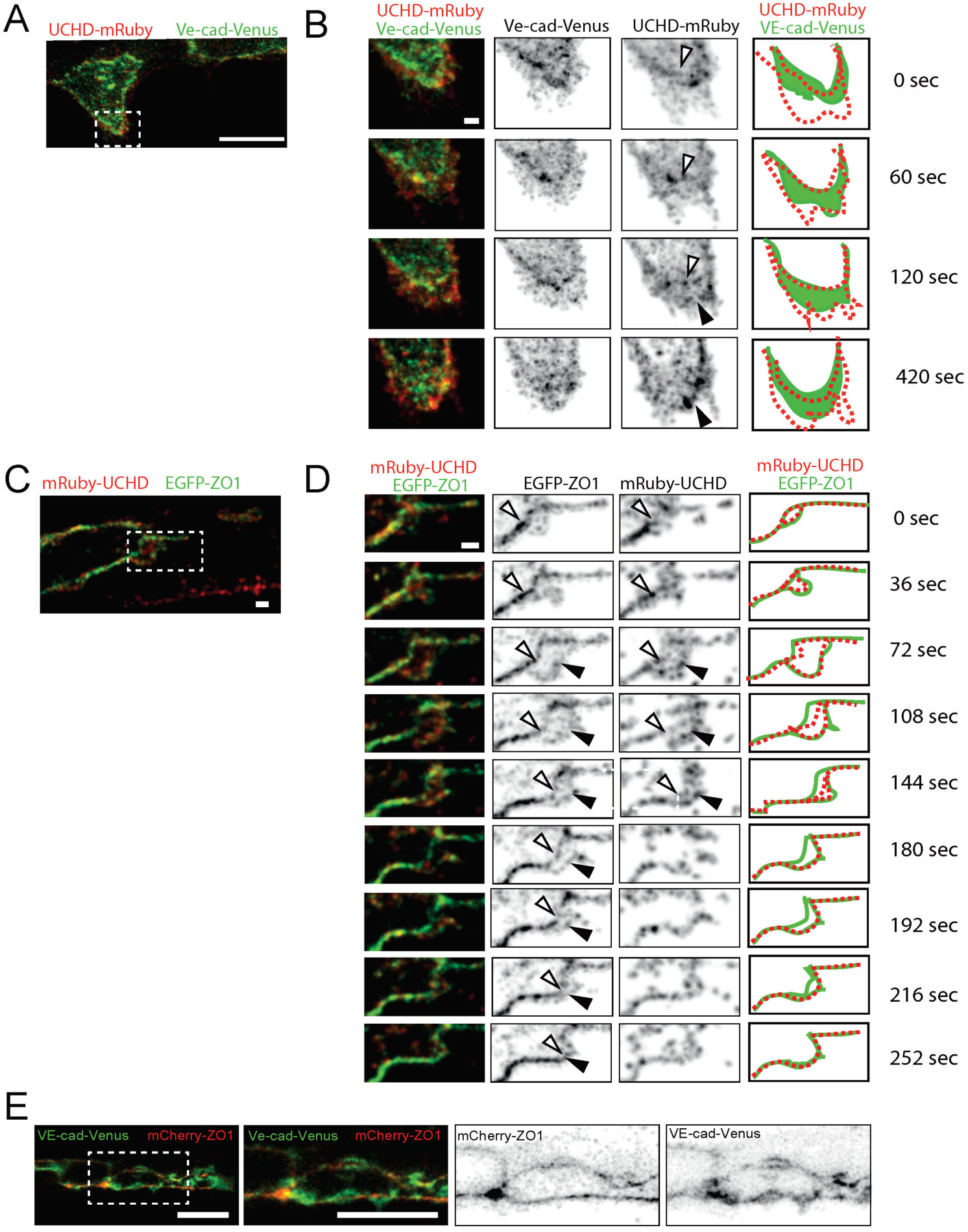
Distinct dynamics of VE-cadherin, F-actin and ZO-1 during JBL formation. A-B) Still images (Supplemental movie 5) of an embryo showing the DLAV around 32 hpf in a embryo expressing both mRuby2-UCHD and VE-cad-Venus *Tg(fli1a:Gal4ff*^*ubs3*^; *UAS:mRuby2-UCHD*^*ubs20*^; *BAC(cdh5:cdh5-Venus))*. B) is a time series magnification of the inset in A). Individual channels are shown in inversed contrast. Similar observation were made in 11 movies. Open arrow head points to established junctions and black arrowhead to pioneering junction. C and D) Still images of an embryo showing) DLAV around 32 hpf (Supplemental movie 5) in an embryo expressing EGFP-ZO1 and mRuby2-UCHD (*Tg(fli1a:Gal4ff*^*ubs3*^;*uas:mRuby2-UCHD*^*ubs20*^; *uas:EGFP-hZO1*^*mubs5*^). Imaged at rate of 12 sec / stack. Similar observations were made in 9 movies. Open arrow head points to established junctions and black arrowhead to pioneering junction. E) Images of endothelial cells in a VE-cad-Venus expressing embryo injected with mCherry-ZO1 encoding plasmid plasmid (*Tg(BAC(cdh5:cdh5-ts));fli1ep:mCherry-ZO1))* (n=7 embryos). Scale bars 1 µm (B,C,D) and 10 µm (A,E).

To directly differentiate the distribution of VE-cadherin and ZO1, we injected mCherry-ZO1 encoding plasmid into VE-cad-Venus transgenic embryos. The differential localization of both proteins confirmed our previous observations and showed that Zo1 distribution is largely restricted to cell junctions. In contrast, VE-cad was also found within areas outside of these junctions (Fig. 5E). Therefore, the respective distribution of Zo1 and VE-cad represent different aspects of JBL formation and illustrates a stepwise mechanism of JBL function. First, F-actin based JBL emanate from EC junctions, which are maintained. The JBL contains diffusely distributed VE-cad. This population of VE-cad precedes formation of the new junction in front of the JBL and may therefore provide adhesive properties for the JBL prior to formation of the new junction. Interestingly, a gradual movement of the old junction towards the new junction was observed in the EGFP-ZO1 movies (Fig. 5D). This indicates that the proximal junction is not resolved in situ, but is actually pulled forward and eventually merges with the distal junction.

### F-actin polymerization and remodeling are required for JBL formation and junction elongation

To elucidate the molecular mechanism underlying JBL function during endothelial cell movements, we examined the requirement of F-actin dynamics by pharmacological interference. Latrunculin B and NSC23766 (a Rac-1 inhibitor) are potent inhibitors of F-actin polymerization and lamellipodial F-actin remodeling, respectively ^18,19^. We used acute treatments to avoid secondary effects and performed live-imaging on rearranging endothelial cell junctions. Inhibition of F-actin polymerization led to pronounced defects in JBL formation. In 5 movies, we observed only 13 JBL, compared to 50 JBL in control embryos (Figure 6A, B). Moreover, those JBL, which did form in the presence of Latrunculin B, lasted longer, indicating additional defects in lamellipodial dynamics. Inhibition of Rac-1 did not interfere as strongly with JBL formation as inhibition of F-actin polymerization. Here, we observed 49 JBL in 10 movies. However, these JBL displayed prolonged duration indicating defects in JBL dynamics (Figure 6A, B). To test whether interfering with F-actin and JBL dynamics has consequences for endothelial cell rearrangements, we analyzed the effect of Latrunculin B and NSC23766 on junction elongation. Both compounds inhibited the elongation of the junction during DLAV formation (Fig. 6C and D) indicating that proper function of JBL is necessary for junctional elongation during anastomosis.

**Figure 6.**
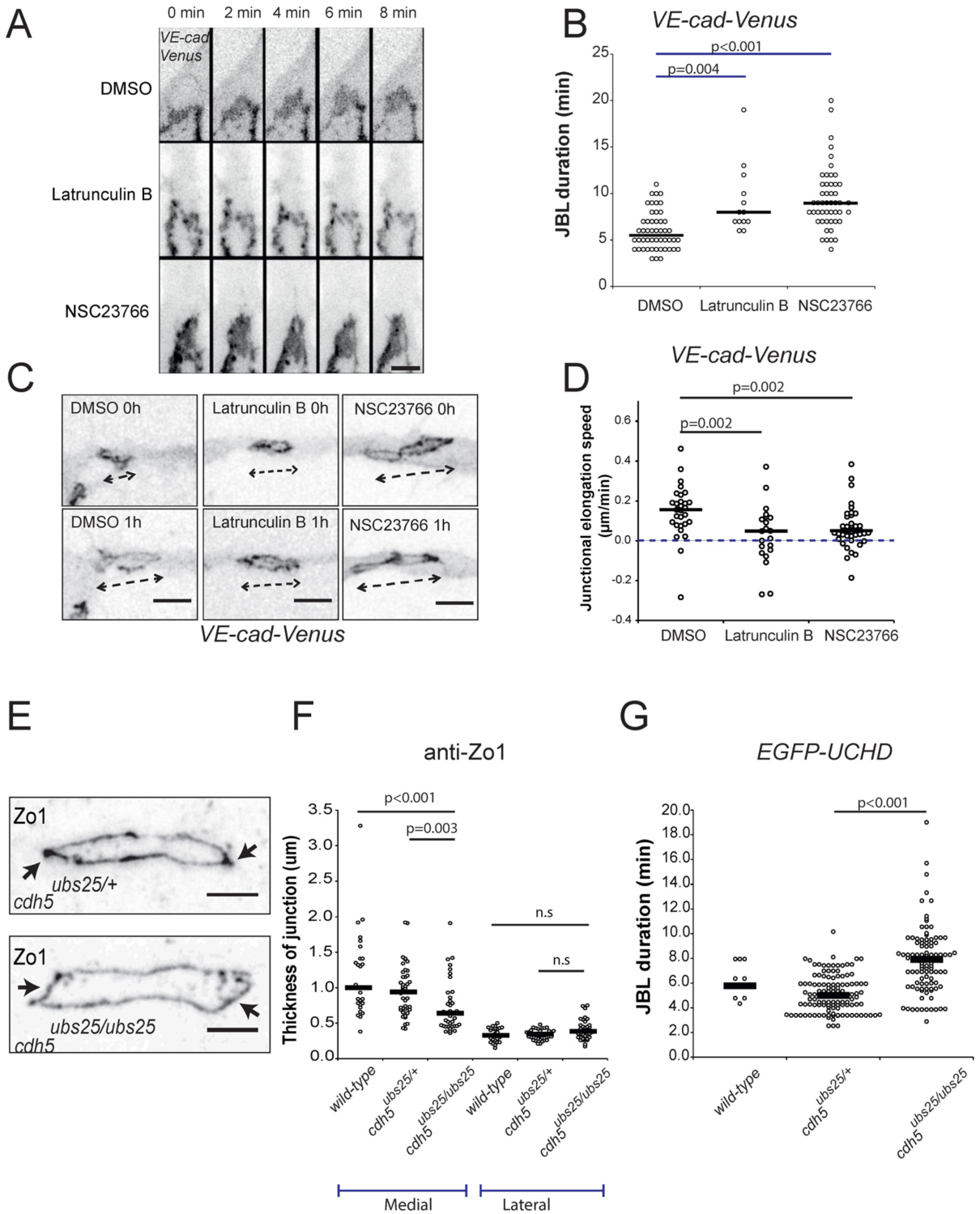
Junction elongation and JBL formation are functionally linked. A) Still images from a movie of an VE-cad-Venus expressing embryo (*Tg*(*BAC*(*cdh5:cdh5-ts*))) during anastomosis in the DLAV (around 32hpf), in the presence of with DMSO (1%), Latrunculin B (150ng/ml) or NSC23766 (900µM). B) Scatter plot quantitation of JBL duration. DMSO, n=50 (6 movies); Latrunculin B, n=13 (5 movies); NSC23766, n=49 (10 movies); black lines show median values. Non-parametric Kruskal-Wallis statistical test was used. C) Confocal images of an *Tg*(*BAC*(*cdh5:cdh5-ts*)) embryo during junctional elongation after DLAV anastomosis. Top panels t=0 and bottom panels after 1h incubation. D) Quantitation of the junctional elongation velocity in the presence of different chemicals using *Tg*(*BAC*(*cdh5:cdh5-ts*)) embryos. DMSO (1%), n=29 junctions (11 embryos); Latrunculin B (150ng/ml), n=21 (6 embryos); NSC23766 (300µM), n=41 (11 embryos). Blue dotted line indicated no movement observed, black lines are medians. Non-parametric Kruskal-Wallis statistical test was used. E) Images of anti-ZO1 immunostained junctions in *cdh5*^*ubs25/+*^ and *cdh5*^*ubs25/ubs25*^ embryos. Arrows point to medial site of the junction. F) Quantitation of the medial and lateral junctional thickness, based on immunostaining for ZO1; *cdh5*^*ubs25/+*^, n=44 junctions (17 embryos); *cdh5*^*ubs25/ubs25*^, n=40 (11 embryos); wild-type n=28 (9 embryos). Black lines are medians. Non-parametric Kruskal-Wallis statistical test was used. G) Quantitation of the duration of JBL based on EGFP-UCHD signal; *cdh5*^*ubs25/+*^, n= 122 (8 embryos), *cdh5*^*ubs25/ubs25*^ n= 103 (8 embryos) and wild-type n=11 (2 embryos). All embryos are *Tg(fli1a:Gal4FF*^*ubs3*^;*UAS.GFP-UCHD*^*ubs18*^). Scalebars: A: 5µm, C: 10µm, E: 5µm.

### VE-cadherin/F-actin interaction is required for JBL function

Next, we wanted to explore the role of VE-cad in JBL function, because several lines of evidence suggested that VE-cad may play an important role in this process. Previously, we have shown that *ve-cad (cdh5* gene in zebrafish) null mutants have defects in endothelial cell and junction elongation during sprout outgrowth and that this phenotype can be copied by the inhibition of F-actin polymerization ^13^. Furthermore, our findings that VE-cad accumulates in JBL prior to junction formation suggests a functional interaction between VE-cad and F-actin during junction elongation. To address this possibility, we generated a targeted mutation in *ve-cad* (*ve-cad*^*ubs25*^/*cdh5*^*ubs25*^), which results in deletion of a portion of the cytoplasmic domain of VE-cad including the ß-catenin binding site, which is essential for VE-cad/F-actin interaction (Figure S3). In homozygosity, this truncation leads to vascular defects similar to those of *ve-cad* null mutant, albeit these phenotypes were somewhat milder (Figure S3 and data not shown). In particular, the dorsal aorta was formed quite normally and we observed blood flow in the DLAV in a subset of *ve-cad*^*ubs25*^ embryos (data not shown). This suggests that the extracellular domain of VE-cad is sufficient to mediate some interendothelial adhesion, which leads to a hypomorphic phenotype. Nevertheless, with respect to SeA formation, we observed the same phenotypes as previously reported for null mutants ^13^, including tip/stalk cell dissociation and junctional gaps (Figure S3E-N and data not shown).

To assess the requirement for VE-cad/F-actin interaction for JBL formation, we first examined whether the junctional rings of *ve-cad*^*ubs25*^ mutants displayed polarized thickness during anastomosis. Immunofluorescent staining for ZO-1 revealed that medial junctions of mutants were narrower compared to heterozygotes, while the thickness of the lateral sides was not affected (Fig. 6E, F). We then tested whether the dynamics of F-actin protrusions are affected in *ve-cad*^*ubs25*^ mutants. Time-lapse analyses in embryos expressing EGFP-UCHD indicated that F-actin protrusions oscillated more slowly in *ve-cad*^*ubs25*^ mutants compared to heterozygotes in a manner similar to how protrusions behaved in the presence of the Rac-1 inhibitor (Figure 6G). Taken together, these findings show that VE-cad plays an important role in F-actin dynamics and that the VE-cad/F-actin interaction is essential for JBL function and junction elongation.

## Discussion

In this study, we have investigated the mechanisms by which junctional dynamics contribute to endothelial cell movements during blood vessel formation *in vivo*. By time-lapse imaging of different structural components of endothelial cell junctions we observe a dynamic and differential deployment of these proteins during junctional remodeling, which leads to the formation of transient lamellipodia-like protrusions, we call junction-based lamellipodia (JBL). Together with our analyses of F-actin dynamics and VE-cad function, our findings suggest a mechanism of cellular and molecular interactions, which allows endothelial cells to use each other as adhesive substrates and for force transmission during cell migration and elongation (Figure 7). In essence, JBL act by ratchet-like mechanism, which - in essence - consists of F-actin based protrusions and VE-cad based interendothelial cell adhesion. While F-actin protrusions provide the motive force, VE-cad based adhesion serves as an intercellular clutch.

**Figure 7.**
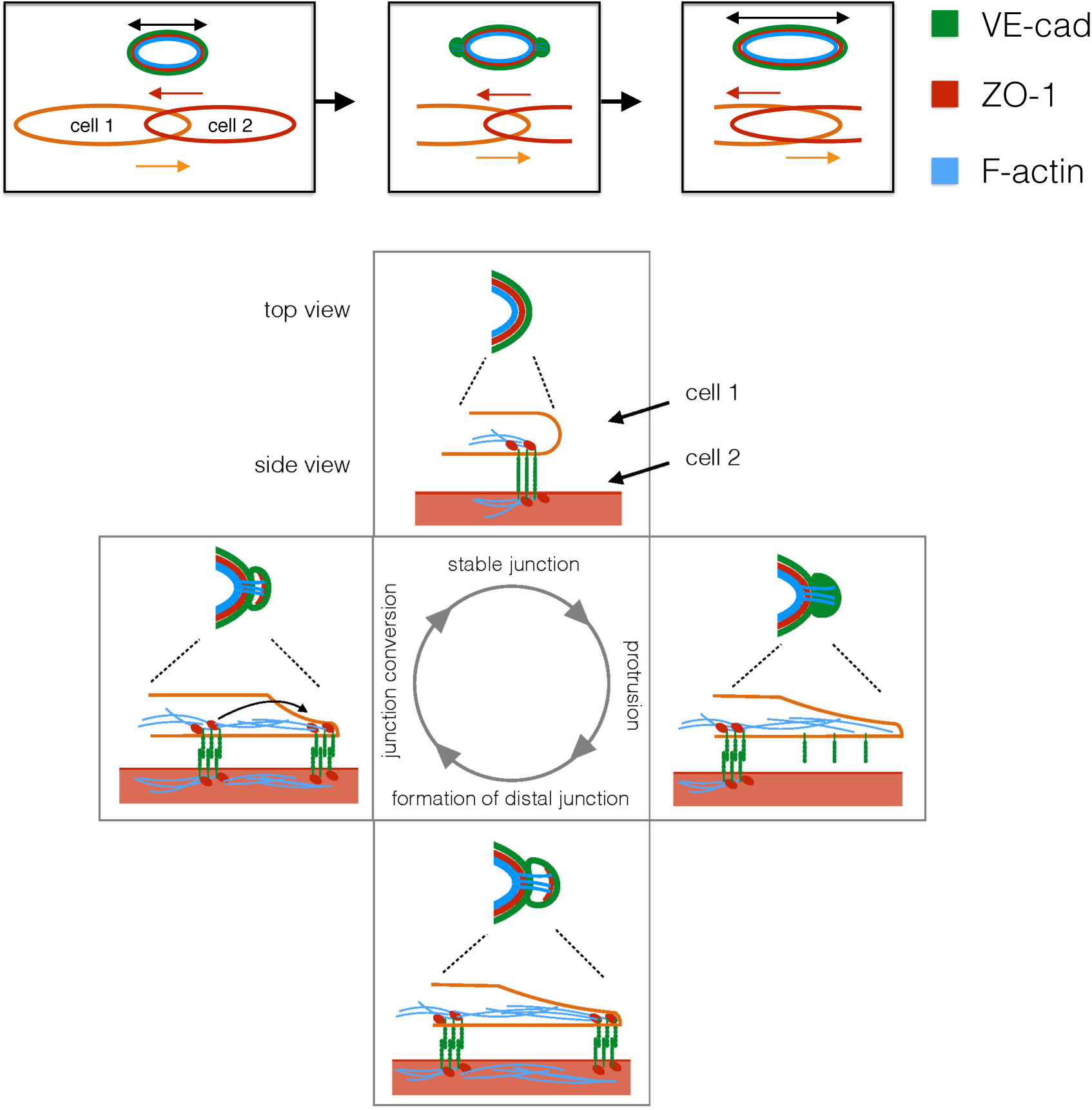
A ratchet-like molecular mechanism of junction remodeling. Top panel: Elongation of an endothelial cell junction during anastomosis. As two endothelial cells move toward and over each other (bottom), the junction becomes elongated. The three proteins investigated in this study are indicated in different colors. Bottom: The proposed oscillatory mechanism of JBL function. Actin protrusions emanate distally from a stable junction. These protrusions also contain diffuse VE-cadherin, but not ZO-1. At the distal end of the protrusion, F-actin, ZO-1 and VE-cadherin become recruited and form a new junction at the front of the JBL. Eventually, dynamic F-actin remodeling (or actin-myosin contractility) pulls the proximal junction towards the new junction.

Our time-lapse experiments show that junction elongation is associated with oscillating JBL, which occur at a frequency of about one every six minutes. This oscillatory behavior, their polarized localization at the leading edge of the junction, their role in cell movements together with their dependency on F-actin polymerization as well as Rac-1 GTPase activity indicates that these protrusions share a functional basis with “classical” lamellipodia. However, and in contrast with “classical” lamellipodia, the protrusions, which we are describing, emanate from interendothelial cell junctions and require VE-cad for adhesion and force transmission.

Several studies have emphasized the importance for VE-cadherin in dynamic endothelial cell interactions including cell rearrangements ^11,12^ and cell elongation ^13^. Here, we show that VE-cadherin can actively contribute to these cell movements via its interaction with the F-actin cytoskeleton. Our model suggests that VE-cadherin provides an extracellular clutch, by generating an intercellular adhesion patch, which serves as a counterfort for intracellular actomyosin contractions. This cell-cell interaction may be analogous to integrin-ECM based adhesion patches of classical lamellipodia. Furthermore, these VE-cadherin adhesion complexes give rise to a distal junction with an underlying F-actin arc. Similar F-actin arcs have been described to be essential for the function of lamellipodia during migration of human endothelial cells (HUVECs) *in vitro* ^20^. Future studies will aim to uncover the exact mechanisms of traction force generation and transmission during JBL driven junction elongation. While F-actin dynamics are essential for junction elongation, we have never observed prominent stress fibers during in this process, suggesting that F-actin based traction forces are acting locally rather than over the longitudinal extent of the endothelial cells.

Oscillatory junctional protrusions of endothelial cells have also been described in HUVECs ^21^. Here, so-called junction-associated intermittent lamellipodia (JAIL) form in similar intervals as JBL. However, several characteristic differences, suggest that JAIL and JBL represent different cellular activities. In contrast to JBL, JAIL formation is preceded by the dissolution of the existing junction. In addition, JAIL do not seem to be associated with cell movements or cell shape changes. Although we have focused our studies on JBL formation and function in the process of blood vessel anastomosis, we observed JBL also within larger caliber vessels such as the dorsal aorta at stages, when endothelial cells are extensively rearranging and undergoing cell shape changes (Lagendijk et al., *in revision*). Our studies therefore indicate that endothelial cells employ JBL as a general means for rearrangements and shape changes during blood vessel assembly and vascular remodeling. Using interendothelial adhesion for force transmission allows dynamic endothelial activities while maintaining the vascular seal. We therefore envision that JBL may underlie many morphogenetic endothelial cell behaviors during blood vessel expansion, normalization, regression and endothelial shear stress response. Remodeling and reorganization of adherens junctions is essential for developmental morphogenesis ^22,23^. Whether similar JBL occur also in different tissues, besides vasculature and endothelial cells, remains to be elucidated.

## Methods

### Fish strains and maintenance

Maintenance of fish and experimental procedures were carried out at the Biozentrum/Universität Basel according to Swiss national guidelines of animal experimentation (TSchV). Zebrafish lines were generated and maintained under licenses 1014H and 1014G1 issued by the Veterinäramt-Basel-Stadt. Fish strains carrying following transgenes and mutations were used in this study: *kdrl:EGFP*^*s843*^ ^24^, *VE-cad-Venus* (*BAC(cdh5:cdh5-TS)*, Anne Karine Lagendijk and Ben Hogan, unpublished), *fli1a:GFF*^*ubs3*^ and *UAS:EGFP-hZO1*^*ubs5*^ ^25^, *UAS:EGFP-UCHDP*^*ubs18*^ ^13^, *UAS:mRuby2-UCHD*^*ubs20*^ (this study), *UAS:mCLAV2-UCHD*^*ubs27*^ (this study), *UAS:mRFP* ^26^ and *cdh5*^*ubs25*^ *(this study)* The fish were maintained using standard procedures and embryos obtained via natural spawning ^27^.

### Generation of Tg(UAS:mRuby2-UCHD)^ubs20^ and Tg(UAS:mClav2-UCHD)^ubs27^

The EGFP sequence of pT24xnrUAS:EGFP-UCHD ^13^ was replaced by the sequence of mRuby2 (amplified from pcDNA3-mRuby2 was a gift from Michael Lin; Addgene plasmid #40260) ^28^ or by the sequence of mClav2 (amplified from pmClavGR2-NT; Allele Biotechnology) to generate the final plasmids pT24xnrUAS:mRuby2-UCHD and pT24xnrUAS:mClav2-UCHD respectively. These final plasmids were injected individually, together with *tol2* RNA into Tg(*fli1a:gal4ff*)^*ubs3*^ embryos.

### Transient expression of mCherry-ZO1 in zebrafish embryos

To transiently express mCherry-Zo1 in endothelial cells of zebrafish embryo, approximately 50 pg of plasmid *fli1ep:mCherry-ZO1* ^29^ was injected together with Tol2-transposase mRNA into 1-4 cell stage embryos (Tg(*BAC*(*cdh5:cdh5-ts*))). 24 hours after injection healthy embryos expressing mCherry were selected, mounted in low-melting point agarose and imaged using Leica SP5 confocal microscope.

### Generation of ve-cadherin mutants

The ve-cadherin truncation allele (*cdh5*^*ubs25*^) was generated using CRISPR/CAS technology (Gagnon et al., 2014). We sequenced exon 12 of *cdh5* (encoding the cytoplasmic domain) of ABC, Tubingen (TU) and tupfel;long-fin (TL) strains and found a potential target sequence in ABC (5’-GGGACCTGCACTCTATG<u>CCATGG</u>-3’). Target guide RNA and Cas9 protein were synthetized by standard procedures ^30^ and co-injected into ABC/TU embryos. Offspring of G0 fish containing germline mutations were screened by PCR analysis for the loss of a *NcoI* restriction site, which is present on the wild-type allele. For subsequent genotyping, multiplex PCR was performed using allele-specific primers:

VE-cad-fwd: 5´-GAAACCCATATCAAACAGACCTGC-3´,
VE-cad-rev: 5´-CAGAGCCGTCTACTCCATAAAGC-3´,
VE-cad-ubs25-fwd: 5´-GACCTGCACTCTATGGAA-3´,
VE-cad-wild-type-rev: 5´-GCAGGAGGTTTCTTTACC-3´.

### Live imaging of zebrafish embryos

Embryos were anesthesized using Tricaine (MS-222, 160mg/l, #E10521 Sigma-Aldrich) and embedded in 0.7% low-melting point agarose (Sigma-Aldrich) supplemented with Tricaine in glass-bottom dish. After the agarose solidified, it was overlaid with E3-medium supplemented with Tricaine. All the imaging was performed at the 28.5**°**C. The imaging was performed using Leica SP5 Matrix confocal microscope equipped with resonance scanner using 63x NA1.2 or 40x NA1.1 water immersion objectives. For imaging of JBL, the time points were 60-120 second intervals and in case of double-color imaging, 12 - 60 seconds.

For the pharmacological experiments, the treatment of embryos with inhibitors (DMSO 1%, Latrunculin B (150ng/ml), NSC23766 (300-900uM)) begun one hour prior to embedding into low-melting point-agarose and confocal imaging. The inhibitors were present throughout the whole experiment.

### Generation of polyclonal rat anti-zf-VE-CAD antibodies

A cDNA fragment encoding a polypeptide comprising the extracellular domain of zebrafish VE-cad (Ala22 to Lys464) was expressed in E. coli using the T7 expression system. The protein was purified on Ni-charged IMAC resin (BioRad) under denaturing conditions. Antiserum was raised in rats by ThermoFisher Scientific using standard immunization procedures.

### Immunofluorescence analysis

Embryos were fixed with 2% paraformaldehyde (Electron Microscopy Sciences) in PBST (PBS+ 0.1% Tween-20) at room temperature for 90 minutes, and immunostained using following protocol: Fixation with 2% PFA/PBST (PBS+0.1% Tween-20) for 90 min at room temperature followed by washes with PBST. After permeabilization (PBST + 0.5% Triton X-100, 30min), the samples were blocked (PBST + 0.1% Triton X100 + 10% normal goat serum + 1%BSA + 0.01% Sodium Azide, overnight, 4°C). Subsequently, primary antibodies were added (diluted in Pierce Immunostain enhancer, #46644, Themofisher Scientific) and incubated overnight at 4°C. After several washes with PBST at room temperature, the secondary antibodies were added (1:2000 dilution in Pierce staining Enhancer) and incubated overnight at 4°C. After several washes with PBST at room temperature, the embryos were mounted onto glass-bottom dishes using low-melting point agarose.

Mouse anti-humanZO-1 (Thermofisher Scientific; used as in ^31^, rat anti-zf-VE-cad and rabbit anti-VE-cad primary antibodies were used. Rat anti-VE-cad was validated in by immunofluorescence by using *ve-cad* null mutant (*cdh5*^*ubs8*^, *Tg(kdlr:EGFP*^*s843*^)) embryos ^13^ as control for specificity (Supplemental information, Fig S2). Rabbit anti-zf-VE-cad has been described and validated earlier ^31^ and also validated with *ve-cad* null mutants earlier (Sauteur et al., 2014). Fluorescent secondary antibodies Alexa-568 goat anti-mouse IgG, Alexa-633 goat anti-rat IgG and Alexa-633 goat anti-rabbit IgG (Thermofisher Scientific) were used.

### Photoconversion experiment

24hpf zebrafish embryos *(Tg(fli1a:GFF*^*ubs3*^*;UAS:mClav2-UCHD^ubs27^)* were embedded in 0.7% low-melting point agarose onto 35mm glass bottom dishes. Some of the tip cells of vascular sprouts of intersegmental arteries were photoconverted using Leica SP5 confocal microscope using 40x NA1.1 water immersion objective. Photoconversion was performed using 405nm laser (20% power) until no obvious increase in converted UCHD-mClav2 signal was observed (conversion time 10-30sec). After this the embryos were allowed to develop for approximately 4 hours before imaging of anastomosis events in dorsal longitunal anastomosing vessel (DLAV).

### Junction elongation experiment

Junctional elongation was analyzed by observing anastomosis and elongation of isolated junctional ring during DLAV anastomosis. Inhibitor treatments Latrunculin B (150ng/ml), NSC23766 (300-900uM) or DMSO (1%) were applied 1hr before mounting of embryos into 0.7% low-melting point agarose and imaging the junctions for 1-2 hours using Leica Sp5 (40x NA1.1 water immersion objective).

### Image analysis and preparation

Image analysis and measurements were performed using FIJI. Deconvolution was performed using Huygens Remote Manager software ^32^. Maximum Z-projections were used. Noise was reduced using Gaussian filtering (radius 1.0) and background subtracted (rolling ball radius 50) using FIJI. Contrast and brightness of images were linearly adjusted. Kymographs were generated from the sum Z-projections of time-lapse series using FIJI. Perpendicular straight line across the junction was drawn and kymograph generated using reslice tool. Publication figures were prepared using FIJI, OMERO figure and Adobe Illustrator.

### Statistical analyses

Statistical analyses were performed using Microsoft Excel and IBM SPSS statistics 22 software. Non-parametric two-sided Kruskal-Wallis H-test, non-parametric one-sample Wilcoxon signed rank test and binomial probability (photoconversion experiments, test probability 0.5) were used. The data reasonably met the assumptions of the tests. In Fig. 1E and F, where the data was non-normal and heteroscedastic similar p-values were obtained with Kruskal-Wallis H-test, Welch´s t-test and median test. No statistical power analysis was used to determine samples size. Systematic randomization was not used. Experiments with *cdh5*^*ubs25*^ (Fig. 6E-G) were performed essentially blinded as the genotype was determined after data capture and analysis. In all other experiments blinding was not used. Samples of low technical quality were excluded from the subsequent analyses. In all figures (exception Fig 2F) the individual data points are plotted and median indicated with horizontal line. In Fig 2F, the data is binned and the number of events in the given bin is plotted.

### Data availability

The data that support the findings of this study are available from the corresponding authors upon reasonable request.

## Acknowledgments

We thank Kumuthini Kulendra for fish care and the Imaging Core Facility of the Biozentrum for microscopy support.

This work has been supported by the Kantons Basel-Stadt and Basel-Land and by a grant from the Swiss National Science Foundation to M.A.. I.P. was supported by a post-doctoral fellowship from the Finnish Cultural Foundation and Foundations’ Post-Doc Pool. M.L. and L.S. by Fellowship of Excellence, Biozentrum, University of Basel.

---

**Supplemental Figure 1.**
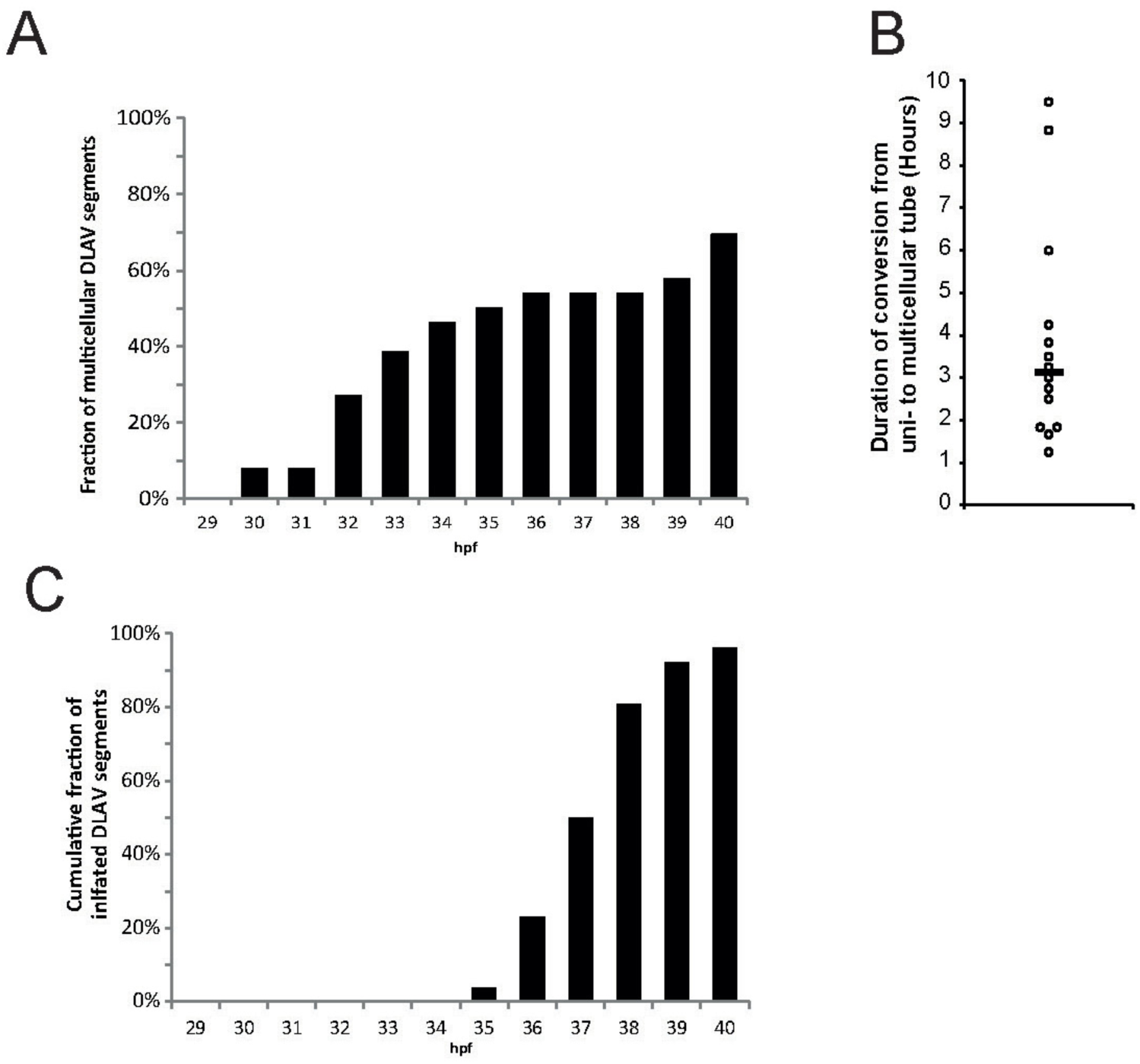
Description of the transition of dorsal longitudinal anastomotic vessel (DLAV) from unicellular to multicellular architecture. A) Quantification of the fraction of multicellular DLAV segments using *Tg(BAC(cdh5:cdh5-ts))* embryos during development. n=26 DLAV segments (8 embryos). B) Quantification of the duration from anastomosis until final conversion into a multicellular tube. Black line is median. n=14 segments (8 embryos). C) Quantification of DLAV segments carrying blood flow during development. n=26 segments (8 embryos). hpf, hours post-fertilization.

**Supplemental Figure 2.**
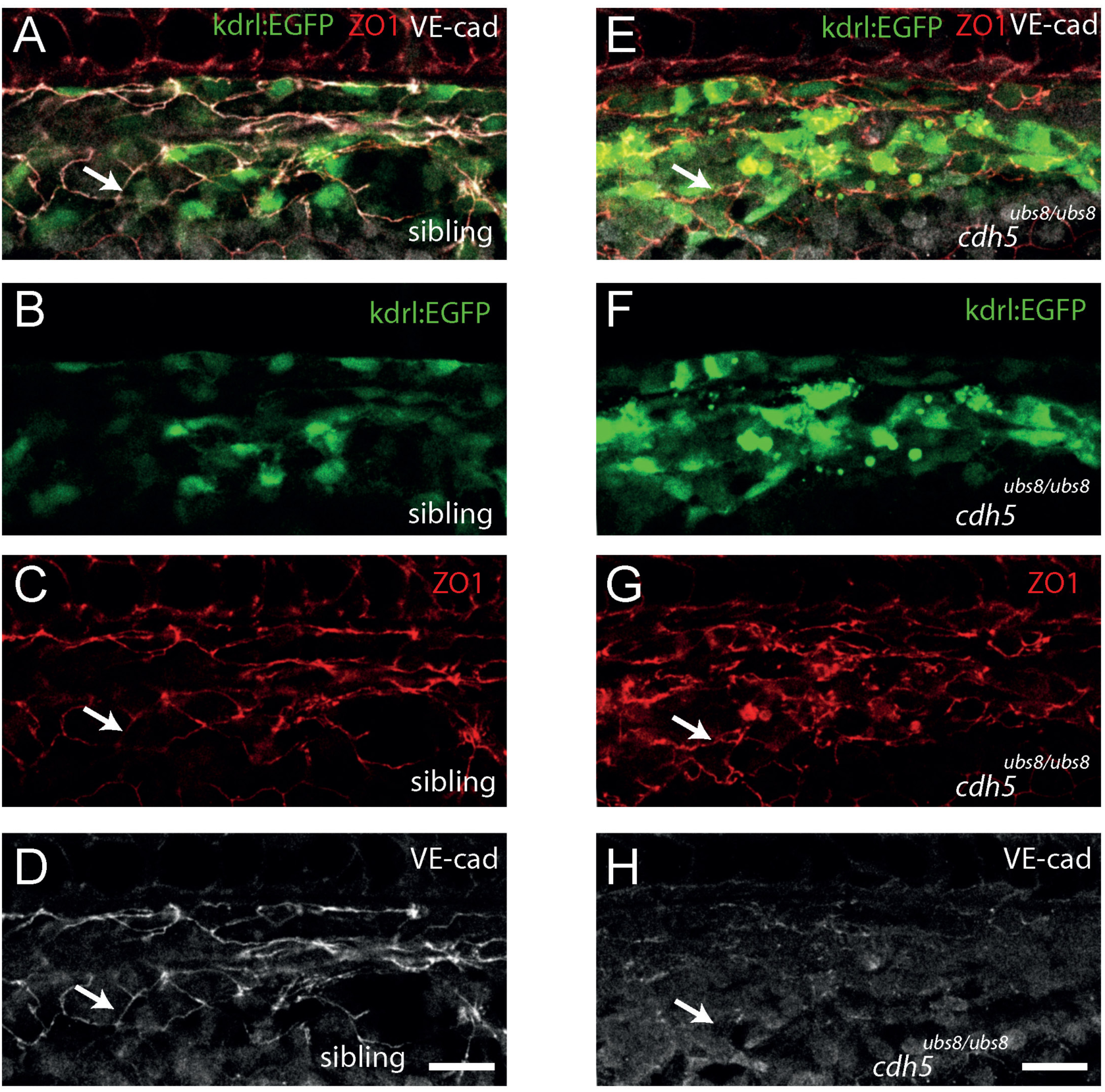
Validation of the rat anti-zf-Ve-cad antibody by whole-mount immunofluorescence analysis. A-H) Confocal images of a *Tg(kdrl:EGFP*^*s843*^) wild-type sibling (A-D) or a VE-cad null mutant embryo (*Tg(kdrl:EGFP*^*s843*^*), cdh5*^*ubs8/ubs8*^) (E-H), stained for VE-cadherin (rat anti-Cdh5) and Zo-1. A and E shows the merged channels, B-D and E-H the individual channels. Arrow points to an endothelial cell-cell junction. Scale bar 20 µm.

**Supplemental Figure 3.**
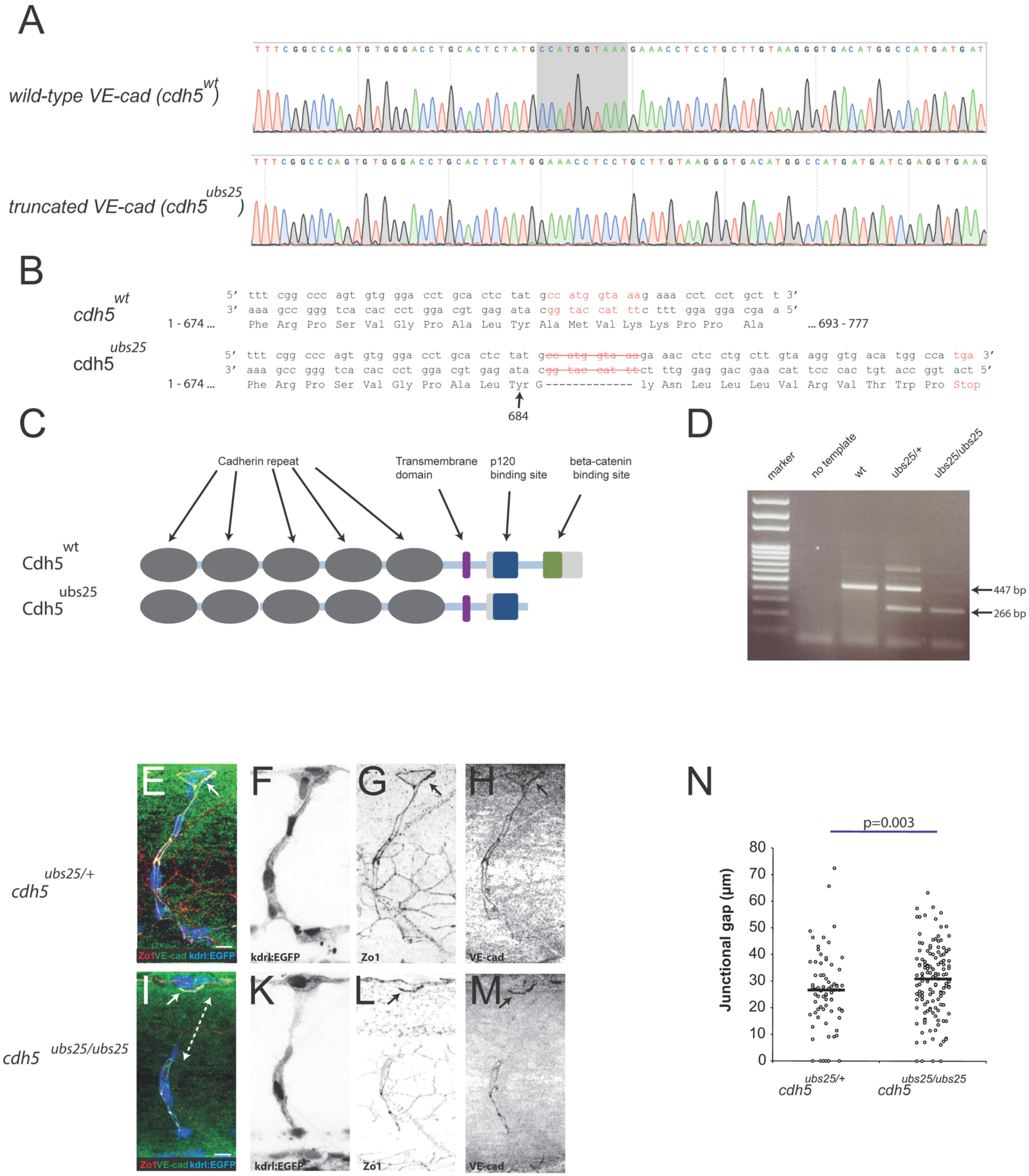
Generation and validation of the ve-cad^*/ubs25*^ (cdh5^*ubs25*^) zebrafish mutant. A) Sequencing chromatogram of wild-type *ve-cadherin* (encoded by *cdh5* gene) sequence (exon 12) and respective truncation mutant *cdh5*^*ubs25*^ sequence. B) The wild-type and mutant DNA sequences and their respective translations. The *ve-cad*^*ubs25*^ mutation leads to a premature stop. C) Schematic illustration of the domains in both full-length wild-type VE-cad and in truncated VE-cad (Cdh5^*ubs25*^). D) Example of genotyping PCR and different *cdh5*^*ubs25*^ allelic combinations. E-M) *Tg(kdrl:EGFP*^*s843*^), *ve-cad*^*ubs25/+*^ (E-H) and *Tg(kdrl:EGFP*^*s843*^), *ve-cad*^*ubs25/ubs25*^ (I-M) embryos stained for VE-cadherin (rabbit antibody, green) and ZO-1 (red). Individual channels are shown in inversed contrast. Both wild-type and mutant VE-cad have junctional localization (solid arrow in panels E,G,I and L). The junctional gap in the VE-cadherin staining of segmental artery of mutant embryo (*cdh5*^*ubs25/ubs25)*^ is marked with dashed double arrow in panel I. N) Quantification of the length of junctional gaps in control (*ve-cad*^*ubs25/+*^, n=72 gaps, 23 embryos) and mutant (*cdh5*^*ubs25/ubs25*^, n=139 gaps, 33 embryos) embryos. Scale bars 10 µm.

## Supplemental Movie Legends

**Supplemental Movie 1.**

Time-lapse imaging of *VE-cad-Venus* (*Tg*(*BAC*(*cdh5:cdh5-ts*))) during DLAV formation starting at 29 hpf, shown in inversed contrast. Time-lapse was imaged at 10min/stack (played at 5 fps, time displayed as hour:min).

**Supplemental Movie 2**

High-resolution time-lapse imaging of *VE-cad-Venus* (*Tg*(*BAC*(*cdh5:cdh5-Venus*))) in the junction of DLAV around 32hpf, shown in inversed contrast. Time-lapse was imaged at 60 sec/stack (played at 5 fps, time displayed as min:sec). Scale bar 5 µm.

**Supplemental Movie 3.**

High-resolution time-lapse imaging of EGFP-UCHD (*Tg*(*fli1a:Gal4ff*^*ubs3*^; *uas:EGFP-UCHD*^*ubs18*^)) in the junction of DLAV around 32 hpf, shown in inversed contrast. Time-lapse was imaged at 90 sec/stack (played at 5 fps, time displayed as min:sec). Scale bar 5 µm.

**Supplemental Movie 4.**

High-resolution time-lapse imaging of EGFP-UCHD (*Tg*(*fli1a:Gal4ff*^*ubs3*^; *uas:EGFP-UCHD*^*ubs18*^)) in the junction of DA around 32hpf, shown in inversed contrast. Time-lapse was imaged at 1min/stack (played at 10 fps, time displayed as min). Scale bar 5 µm.

**Supplemental Movie 5.**

High-resolution time-lapse imaging of mRuby-UCHD (red) and VE-cad-Venus (green) (*Tg(fli1a:Gal4ff*^*ubs3*^; *uas:mRuby2-UCHD*^*ubs20*^; *BAC(cdh5:cdh5-ts))* in the junction of DLAV around 32hpf. The time-lapse was imaged at 30 sec / stack (played at 5 fps, time displayed as min:sec). Scale bar 5 µm.

**Supplemental Movie 6.**

High-resolution time-lapse imaging of mRuby-UCHD (red) and EGFP-ZO1 (green) (*Tg(fli1a:Gal4ff*^*ubs3*^; *uas:mRuby2-UCHD*^*ubs20*^; *uas:EGFP-hZO1*^*ubs5*^)) in the junction of DLAV around 32 hpf. Time-lapse was imaged at 12 sec/stack (played at 5 fps, time displayed as min:sec). Scale bar 1 µm.

